# TLR3 Expression in Villus-Like Enterocytes Drives IFN-III Responses to Enteroviruses

**DOI:** 10.64898/2026.04.01.715830

**Authors:** David N. Hare, Carolyn B. Coyne

**Affiliations:** Department of Integrated Immunobiology, Duke University, Durham, NC, USA; Duke Human Vaccine Institute, Durham, NC

## Abstract

Enteroviruses initiate infection at the intestinal epithelium but can spread systemically to cause severe disease. Although both MDA5 and TLR3 have been implicated in enterovirus sensing, the mechanisms by which the intestinal epithelium detects these viruses remain poorly defined. To address this, we infected human intestinal organoids (enteroids) with echovirus 11 (E11) and compared responses in models differentiated to mimic either crypt-like or villus-like epithelium. Villus-like enteroids produced significantly more type III interferons (IFN-λs) following E11 infection or treatment with the dsRNA mimetic poly I:C, and exhibited heightened responsiveness to IFN-λ signaling. Single-cell RNA sequencing (scRNA-seq) of infected enteroids revealed that E11 broadly infected epithelial cell types, but IFN-λ expression was largely restricted to mature enterocytes. Notably, enterocyte differentiation was also associated with upregulation of innate immune genes. Using CRISPR-Cas9 knockout enteroids, we found that TLR3 signaling was essential for intestinal IFN-λ responses to E11 infection, whereas loss of MAVS, the adaptor for MDA5, had no effect. Together, these data support a model in which mature enterocytes serve as key sensors of enterovirus infection via TLR3, triggering a localized IFN-λ response that may help restrict viral spread.

**Importance:** Enteroviruses are commonly circulating viruses that can cause a broad spectrum of disease, particularly in pediatric populations. Innate immune sensing in the intestinal epithelium likely plays a critical role in determining the outcome of enterovirus infections. Deficiencies in TLR3 signaling, IFN-λ responses, or crypt-villus architecture may contribute to severe disease presentations. Our findings highlight the importance of TLR3-mediated sensing in mature enterocytes, which may also have broader implications for other intestinal viruses, as well as for inflammatory conditions like inflammatory bowel disease. A deeper understanding of how TLR3 sensing and IFN-λ production are regulated in the intestine could inform new therapeutic strategies aimed at modulating mucosal sensitivity to viral nucleic acids and enhancing antiviral defense.

## Introduction

Enteroviruses can cause severe disease in neonates, including meningitis, liver failure, and, in rare cases, even death (1, 2). Many clinically important enteroviruses including poliovirus, coxsackieviruses, enterovirus A71, and echoviruses are transmitted via a fecal-oral route and first encounter the intestinal epithelium during host entry. To cause systemic disease, enteroviruses must breach the intestinal barrier to access secondary sites such as the liver, heart, and central nervous system where they are associated with severe disease.

The small intestinal epithelium is organized into crypts and villi and is constantly replenished by intestinal crypt stem cells (ISC). ISCs that leave the crypt niche differentiate into rapidly dividing transit amplifying cells before becoming enterocytes, BEST4 ionocytes, tuft cells, goblet cells, Paneth cells, or enteroendocrine cells (EEC) (3). Enterocytes comprise the largest proportion of epithelial cells in the small intestine and are involved in nutrient absorption. As enterocytes migrate toward the villus tips, they undergo terminal differentiation and ultimately shed into the intestinal lumen ∼4 days after their generation (3). Because of their rapid turnover, enterocytes are constantly being replenished by ISCs. Maturation along the crypt-villus axis is driven by decreasing levels of Wnt and increasing levels of bone morphogenic protein (BMP) (3, 4). ISCs grown *in vitro* can generate three-dimensional organoids, or enteroids, that can differentiate into nearly all intestinal epithelial cell types (5). To maintain ISC stemness, culture media must include Wnt, BMP inhibitors, and other growth factors; however, these conditions can limit epithelial differentiation and cellular diversity (6). Recent protocols overcome this limitation by supporting robust ISC expansion followed by directed differentiation into crypt-like or villus-like enteroids (4, 6, 7).

The first line of defense against enterovirus infection in the intestine are antiviral genes coordinated by the antiviral cytokine interferon (IFN) (2). The most studied IFN-I genes are IFNB (IFN-β) and 13 IFNA (IFN-α) subtypes. IFN-III genes include IL29 (IFN-λ1), IL28A/B (IFN-λ2/3), and IFNL4 (IFN-λ4) (8). IFN-I signals through the IFN-α/β receptor (IFNAR1 and IFNAR2) while IFN-III signals through the IFN-λ receptor (IFNLR1 and IL10RB) (9–11). Both IFNAR and IFNLR activate a common transcriptional complex and upregulate similar sets of IFN-stimulated genes (ISGs) (12). IFNAR1, IFNAR2, and IL10RB are widely expressed while IFNLR1 is primarily expressed in epithelial cells, including all mucosal epithelia, hepatocytes, and trophoblasts (12–15).

Production of IFN-I and IFN-III is dependent on recognition of molecular patterns like double-stranded RNA (dsRNA) by innate pattern recognition receptors. dsRNA is produced as a replication intermediate of RNA viruses and released into the cytosol or extracellular space (16, 17). TLR3 recognizes dsRNA in endosomes, often taken up from nearby virus-infected cells, and signals through the adaptor protein TRIF (16). MDA5 recognizes intracellular dsRNA in the cytosol of infected cells and signals through the adaptor protein MAVS on mitochondria and peroxisomes (18, 19). Both TLR3-TRIF and MDA5-MAVS signaling converges on activation of IRFs and NF-κB which upregulate IFN-I and IFN-III (20–22). TLR3 and MDA5 are both expressed basally in intestinal epithelial cells but are additionally regulated by IFN (23, 24).

Genome-wide association studies have identified mutations in TLR3 and MDA5 in some children with enterovirus encephalitis, suggesting they play a role in restricting enterovirus infection (25, 26). Studies in mice suggest that the dsRNA sensors TLR3 and MDA5 can sense and restrict replication of several enteroviruses including coxsackievirus B (CVB), coxsackievirus A (CVA), enterovirus A71 (EV-A71), and poliovirus (27–32). However, these studies employed intraperitoneal injection, thereby bypassing the intestinal barrier and limiting the ability to determine which cell types are responsible for sensing echoviruses via TLR3 or MDA5. Enteroviruses preferentially induce IFN-λs in human intestinal enteroids (33, 34). In neonatal mice, ablation of IFNLR impaired the clearance of echovirus specifically from the gastrointestinal tract, highlighting the importance of IFN-λ signaling in mucosal antiviral defense (35). However, the specific epithelial cell types responsible for IFN-λ production and the pattern recognition receptors responsible for this induction remain unknown.

To investigate where and how IFN-λs are produced in response to enterovirus infection, we leveraged recent advances in the expansion and directed differentiation of human intestinal enteroids. These protocols enabled both genetic manipulation and the generation of crypt-like or villus-like enteroids containing nearly all intestinal epithelial cell types. Using this model, we found that echoviruses infect a broad range of epithelial cells, but IFN-λ is produced primarily by mature enterocytes through the TLR3–TRIF–IRF3 signaling axis. We further show that key innate immune genes, including IFNLR1 and TLR3, are enriched in mature enterocytes within villus-like enteroids, consistent with their heightened responsiveness to IFN-λ, double-stranded RNA, and echovirus infection. These results highlight mature enterocytes as key antiviral sentinels in the intestinal epithelium and provide mechanistic insight into how epithelial differentiation governs innate immune responses to enterovirus infection.

## Results

### Modeling Crypt–Villus Architecture Using Patterned Human Intestinal Enteroids

To model intestinal crypts and villi, we designed a patterning and differentiation protocol based on previously published enteroid models (4, 6). Fourteen days in patterning media containing IL-22 promotes Paneth cell differentiation and yields a cellular composition resembling that of the intestinal crypt. (6). Three days in differentiation media lacking Wnt3a, R-Spondin-1, and Noggin (WRN) and containing BMP-2/4 promotes the maturation of enterocytes, goblet cells and enteroendocrine cells and a cellular composition resembling intestinal villi (4). To mimic the architecture of intestinal crypts or villi, we cultured human enteroids in patterning media for 11 days, followed by an additional 3 days in either patterning or differentiation media. Compared to enteroids maintained in expansion media, those subjected to patterning or differentiation protocols developed thicker epithelial walls and irregular morphologies, consistent with the presence of differentiated enterocytes and secretory cell types (**Figure 1A**). Immunofluorescence microscopy for markers of EECs (chromogranin A - CHGA), goblet cells (mucin 2 - MUC2), Paneth cells (lysozyme C - LYZ), transit amplifying cells (KI67), and ionocytes (bestrophin 4 - BEST4) confirmed that these specialized cell types were present in these enteroid models (**Figure 1B**).

**Figure 1.**
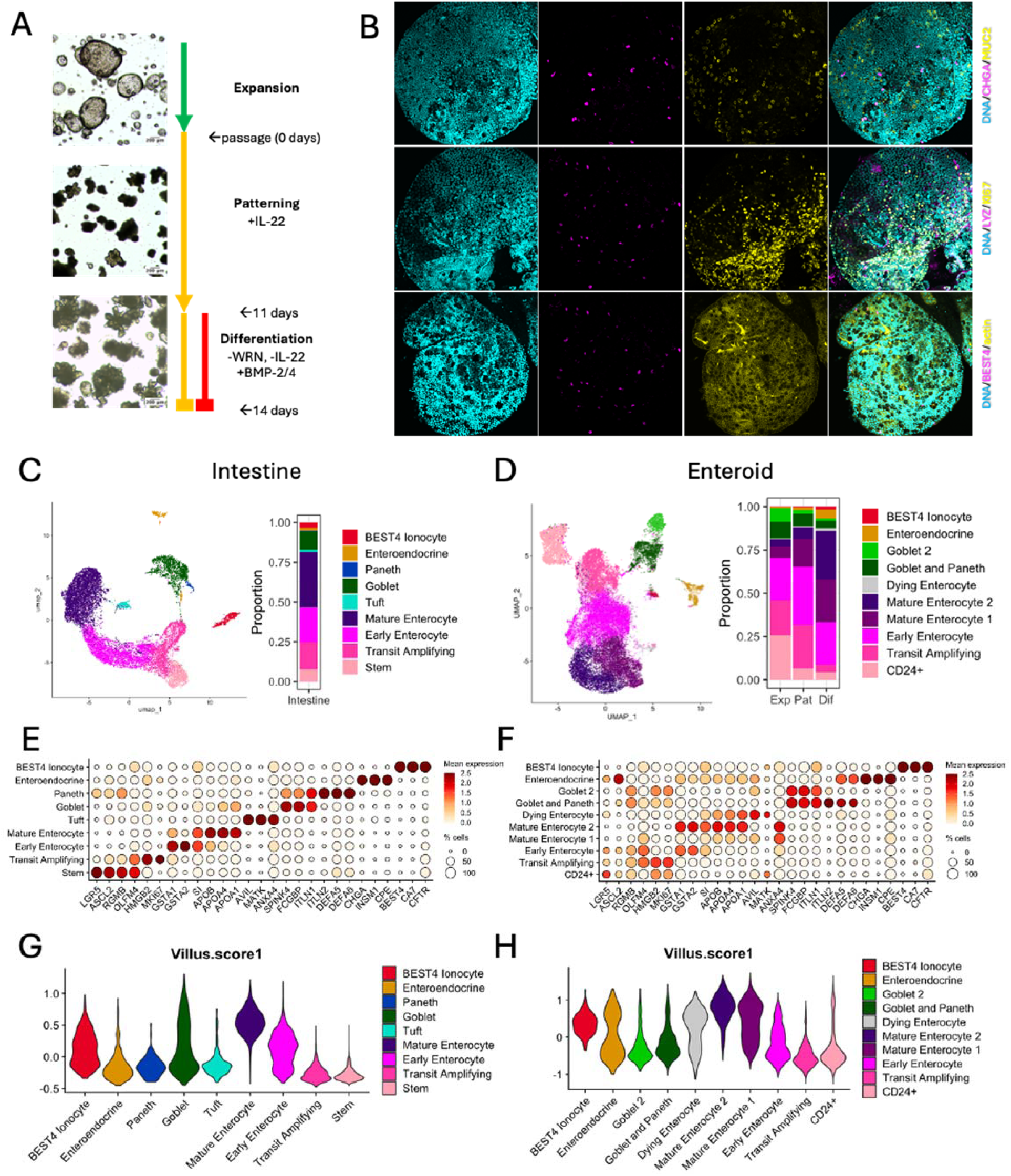
Patterned and differentiated enteroids mimic human intestinal crypts and villi. Human enteroids grown in expansion media are passaged and grown in patterning media for 11 days before 3 days in either patterning or differentiation media to model intestinal crypts or villi. Brightfield images of enteroids after 7 days in expansion, 7 days in patterning, or 11 days in patterning and 3 days in differentiation (A). Patterned enteroids were stained for mucin 2 (MUC2), chromogranin A (CHGA), lysozyme C (LYZ), KI67 and DNA, while differentiated enteroids were stained for bestrophin 4 (BEST4) and actin and visualized using fluorescence microscopy. Markers alone or full composites are shown (B). UMAP and bar plots show distribution and frequency of cell type clusters (C-D). Dotplots show the mean expression and frequency of marker gene expression across cell type clusters (E-F). Violin plots show the distribution of villus scores among cells in each cell type cluster (G-H).

To study the cellular composition of patterned and differentiated enteroids, we grew enteroids using patterning, differentiation, or expansion protocols and performed single-cell RNA sequencing (scRNA-seq). To compare this scRNA-seq dataset to human intestinal epithelial tissue, we pooled human small intestinal scRNA-seq datasets from published studies and analyzed them in parallel (36–38). We used CellRanger to map reads and a modified Seurat pipeline to integrate datasets, perform PCA, and cluster cells. Principle components were used to create UMAP projections of both datasets (**Figure 1C-D**). We used canonical marker genes identified in the human intestinal tissue datasets to validate the cell types present in enteroid models (**Figure 1E-F**). Using these genes, we identified clusters corresponding to the major intestinal epithelial cell types in both intestinal tissue and our enteroids. These included transit amplifying cells, early enterocytes, mature enterocytes, goblet cells, Paneth cells, EECs, and BEST4+ ionocytes (**Figure 1E-F**). Patterned enteroids contained a high proportion of transit amplifying and early enterocytes while differentiated enteroids contained more BEST4+ ionocytes, EECs, and mature enterocytes, as expected (**Figure 1D**). These differences in cell type composition are consistent with observed differences between intestinal crypts and villi (36, 39, 40).

Because enteroids lack a true spatial crypt-villus organization, we assessed whether enterocytes within enteroids follow a similar maturation trajectory by using AddModuleScore to assign each cell a villus score based on expression of established villus-associated genes (41). In both intestinal tissue and enteroids, enterocyte villus scores increased progressively as cells transitioned from transit-amplifying cells to mature enterocytes (**Figure 1G–H**). Together, these results demonstrate that enteroids capture key aspects of enterocyte maturation, supporting their utility as a model of intestinal epithelial biology.

To assess whether intestinal epithelial cell types differ in their capacity to sense and respond to enterovirus infection, we analyzed expression of key viral receptors and innate immune genes using our enteroid and small intestine scRNA-seq datasets. The echovirus receptor FcRn (FCGRT) was broadly expressed across epithelial populations in both systems, suggesting multiple cell types may be permissive (**Figure 2A–B**). In contrast, expression of the echovirus attachment factor DAF (CD55) was more variable, implying potential differences in susceptibility among epithelial subsets for isolates binding this receptor.

**Figure 2.**
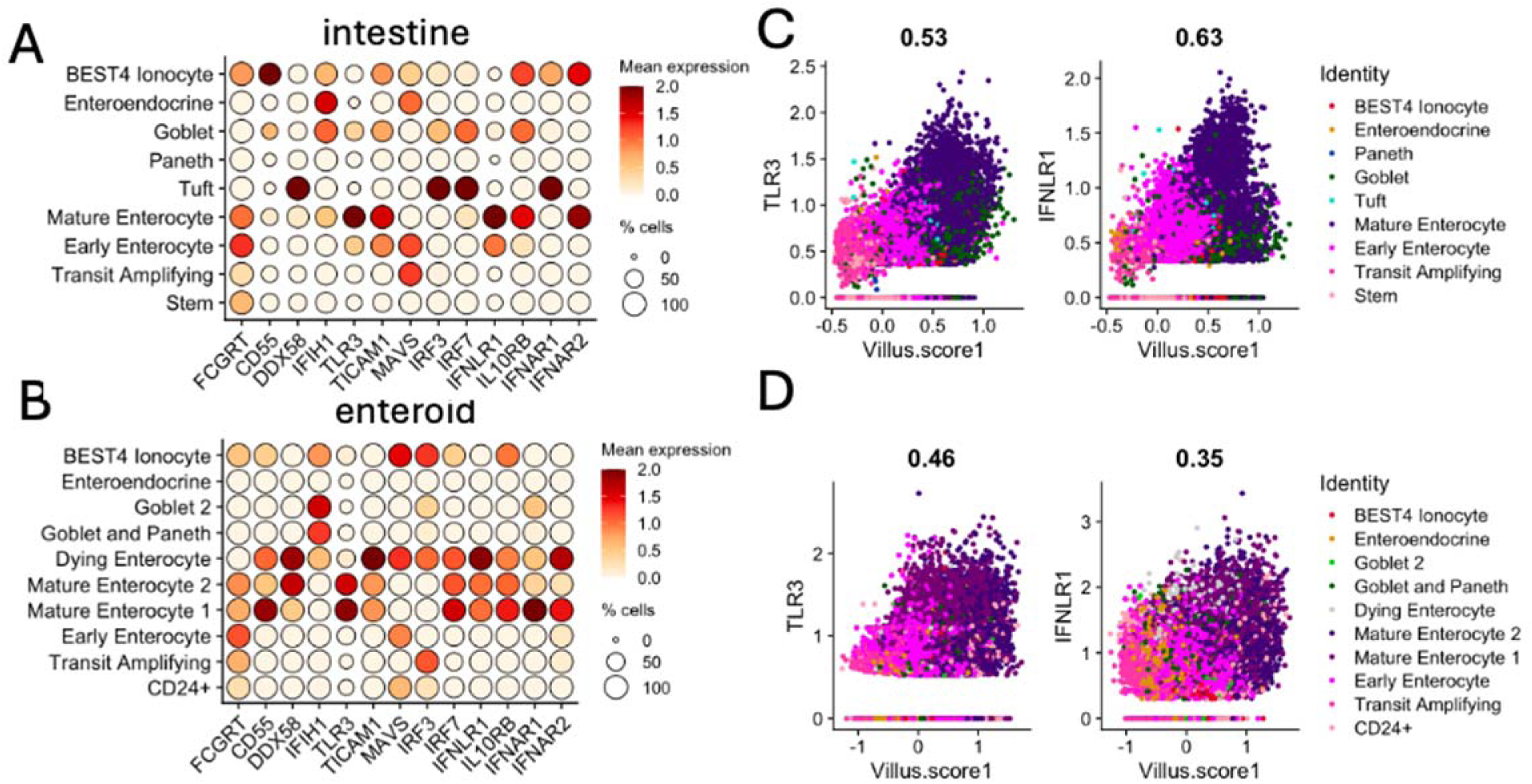
TLR3 and IFNLR1 expression correlates with enterocyte maturity. Dot plots show the mean expression and frequency of genes of interest for intestinal enterovirus-host interactions in both intestine (A) and enteroids (B). Scatter plots show the mean expression of TLR3 and IFNLR with vilus score in intestine (C) and enteroids (D).

We next assessed expression of innate immune genes involved in sensing viral RNA and responding to IFNs. Several of these genes showed elevated basal expression in mature enterocytes, including *TLR3* and *IFNLR1*. Both genes were consistently upregulated in mature enterocytes in enteroids and intestinal tissue. Moreover, their expression correlated positively with villus score in both datasets (**Figure 2C–D**), indicating that as enterocytes differentiate, they increasingly express components of dsRNA sensing and type III interferon signaling pathways. These findings suggest that enterocyte maturation correlates with upregulation of innate immune pathways that could enhance responsiveness to dsRNA and IFN-λ along the villus axis.

### Differentiated Enteroids are More Responsive to IFN-λ, dsRNA, and Echovirus Infection

The ability to model intestinal villi (differentiated) or crypts (patterned) with enteroids enabled us to assess how enterocyte differentiation shapes responsiveness to IFN-IIIs and dsRNA. Treatment of patterned and differentiated enteroids with recombinant IFN-λ1 revealed that several measured ISGs were more strongly upregulated in differentiated enteroids (**Figure 3A**). Next, we treated both patterned and differentiated enteroids with the dsRNA-mimetic poly I:C. Differentiated enteroids exhibited a stronger induction of IFN-λ1 and IFN-λ2 transcripts compared to patterned enteroids (**Figure 3B**). At the protein level, IFN-λ1 production was modestly increased in differentiated enteroids following poly I:C treatment (**Figure 3C**). These data suggest that differentiated enteroids mount a more robust antiviral response to both IFN-III and dsRNA, consistent with elevated expression of innate immune sensors and receptors in mature enterocytes.

**Figure 3.**
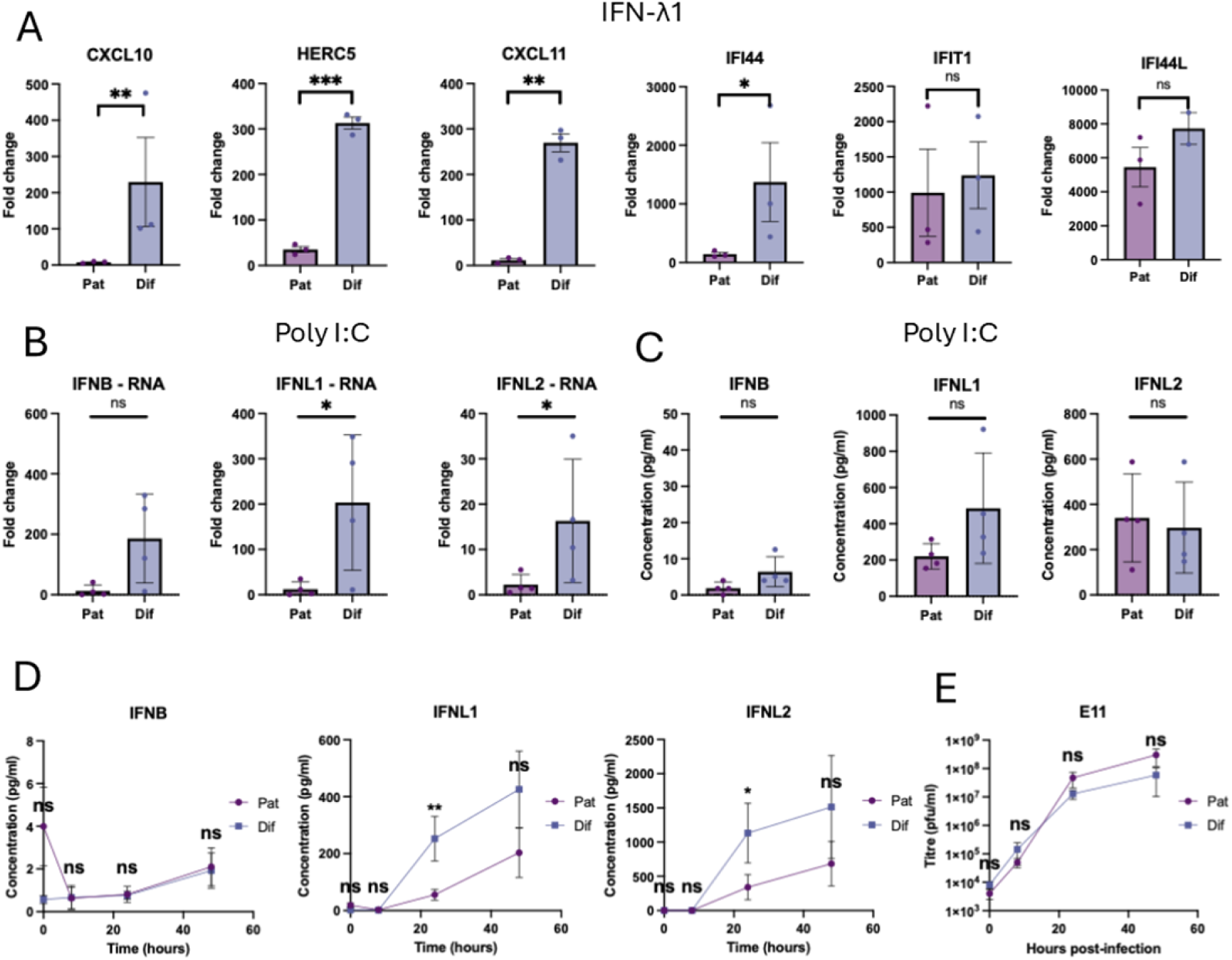
Enteroid differentiation enhances responses to IFN-III and E11. Enteroids grown through patterning (Pat) or differentiation (Dif) protocols were treated with 5 ng/ml IFN-λ1 for 24 h and RNA harvested to measure representative ISGs by qPCR (A). Pat or Dif enteroids were treated with 10 µg/ml poly I:C for 24 h and supernatants and RNA harvested to measure IFNB, IFNL1, and IFNL2 RNA by qPCR (B) and protein by Luminex (C). Pat or Dif enteroids were infected with 1 pfu/cell E11 and 100 µl of supernatant sampled 8, 24, and 48 hours post-infection. IFNB, IFNL1, and IFNL2 protein were measured by Luminex (D) and infectious E11 at each time was measured by plaque assay (E). Mean and standard error are shown for all experiments and the limit of detection (dotted line) is shown for Luminex experiments. Mean values were compared by log normalized t tests and multiple comparisons were corrected for with the Holm-Sidak method. ns: not significant, *: p<0.05, **:p<0.005, ***:p<0.0005.

Previous studies in human enteroids found that echovirus 11 (E11) infected both enterocytes and EECs, was more cytotoxic than CVB or EV71, and was sufficient to induce detectable IFN-III *in vitro* (*33, 34*). To determine whether crypt- or villus-derived intestinal cells are more responsive to E11 infection, we infected both patterned and differentiated enteroids. Supernatants from differentiated enteroids infected with E11 contained significantly more IFN-λ1 and IFN-λ2 than patterned enteroids (**Figure 3D**). Consistent with prior results, no IFN-β was detected in either patterned or differentiated enteroids (**Figure 3D**) (33). Despite the difference in IFN-III production, E11 replication was unchanged between patterned and differentiated enteroids over a 48-hour time course (**Figure 3E**). These results suggest that differentiated enteroids are able to mount a stronger IFN-III response to E11.

### A Subset of Mature Enterocytes Upregulate IFNL Following Echovirus Infection

Given the heightened IFN-λ response of differentiated enteroids to E11, we next examined which cell types harbored E11 genomes and how these cells responded to infection. To do this, we infected differentiated enteroids with E11 and performed scRNA-seq. As expected, differentiated enteroids contained a mixture of enterocytes, goblet cells, and EECs (**Figure 4A-B**). Most clusters were equally distributed between mock and E11 infected samples, except for Dying Enterocytes, WNT9A Enterocytes, and IFNL Enterocytes which were more abundant in the E11 infected samples (**Figure 4A**). In the E11 infected samples, approximately 40% of cells contained E11 genomes (E11+) and enterocytes, goblet cells, and EECs were all similarly susceptible to E11 (**Figure 4C**). Approximately 4% of cells in the infected samples contained IFNL transcripts and were clustered together (**Figure 4D**).

**Figure 4.**
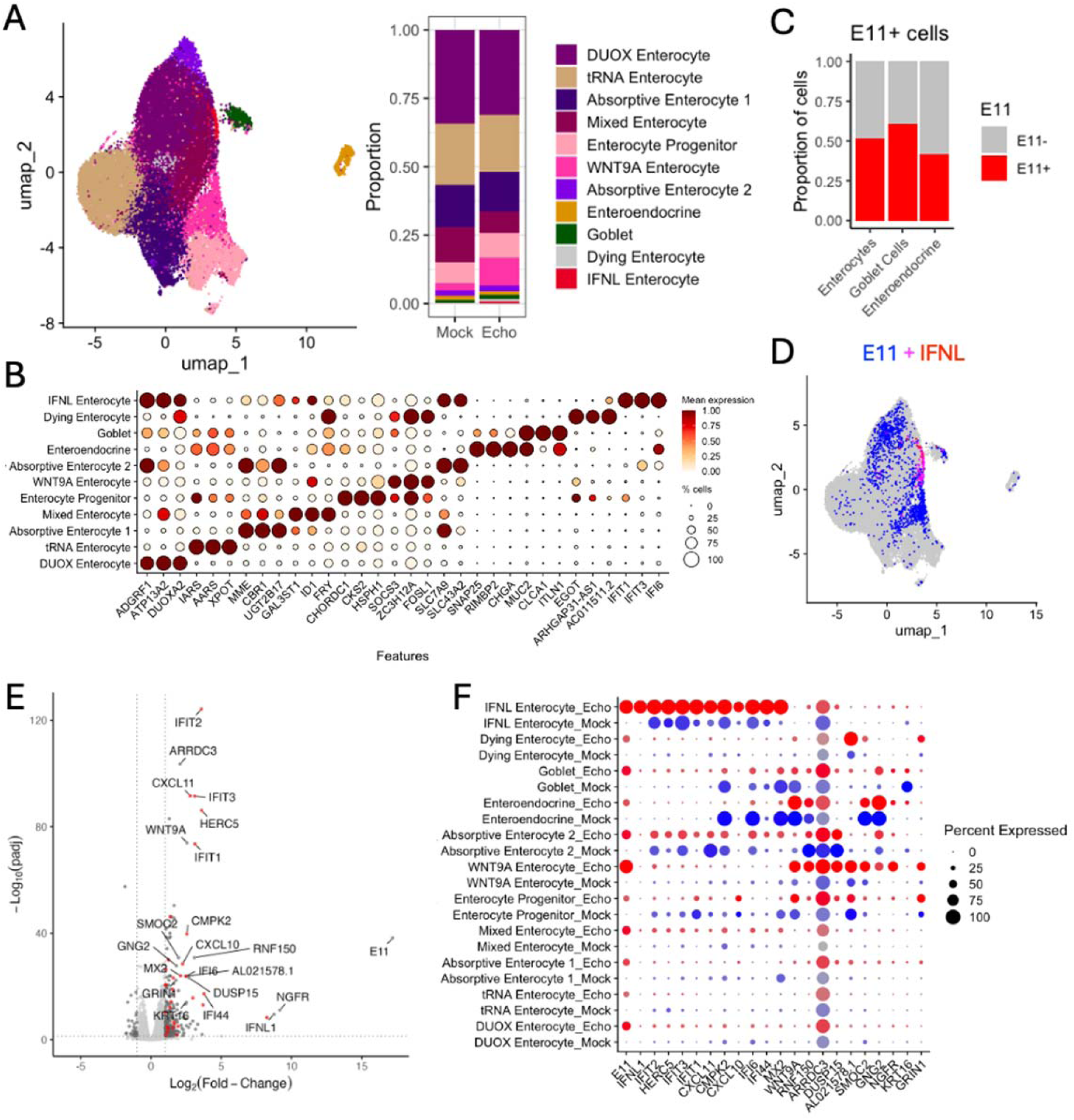
IFN-III is upregulated by a subset of E11-infected cells in differentiated enteroids. UMAP and bar plot from scRNA-seq data show the distribution and frequency of cell type clusters (A). Dot plot shows the mean expression and frequency of top marker genes for each cluster (B). Bat plot of E11+ (red) and E11-(grey) cells shows the frequency of E11+ cells in clusters after amalgamating the enterocyte clusters (C). UMAP with E11 (blue), IFNL score (red), and overlap (magenta) shows the distribution of E11+ and IFNL+ cells (D). Volcano plot shows the -Log_10_(p-value) and Log_2_(fold change) of all genes differentially expressed between E11- and mock-infected enteroids. Pseudobulk analysis was used to calculate fold-change and p-value. Genes with >2 fold change and <0.05 p-value are shown in red (ISGs) and dark grey (non-ISGs) and select genes are labeled (E). Dot plot shows the mean expression and frequency of the top 10 differentially expressed genes classified as ISGs or non-ISGs across all clusters. Clusters were split into E11 infected (red) and mock infected (blue) samples to show changes in expression (F).

We next performed pseudobulk analyses to compare gene expression between infected and uninfected populations. Differentially experessed genes (DEGs) were defined by a greater than 2-fold change and a p-value less than 0.05. We sorted DEGs into ISGs and non-ISGs using an online database of human samples stimulated with IFN-I/III (42). Pseudobulk analysis revealed robust upregulation of IFNL and numerous ISGs, along with many non-ISG transcripts in E11 infected samples (**Figure 4E**). To identify the most differentially expressed genes we ranked DEGs by fold change and p-value and used the sum of those ranks to assign each gene a combined score and identify and label genes with large fold change and low p-value (**Figure 4E**). Of these DEGs, the ISGs and non-ISGs were upregulated in distinct E11 infected clusters (IFNL Enterocyte and WNT9A Enterocyte) (**Figure 4F**). This suggests two distinct responses to infection based on the presence/absence of IFN signaling.

To better examine differential responses to E11 infection, we subsetted and re-clustered all E11+ cells (**Figure 5A**). Using AddModuleScore, we calculated an ISG score and a non-ISG score for each cell based on the top 10 ISGs and top 10 non-ISGs. Non-ISG scores were highest in WNT9A⁺ enterocytes and LGR5⁺ enterocytes, whereas ISG scores were highest in the IFNL⁺ enterocyte population, with minimal overlap between the two groups (**Figure 5B**). We found that the ISG score positively correlated with the villus score of E11⁺ cells, whereas the non-ISG score showed no such relationship (**Figure 5C**). Moreover, E11⁺ cells expressing IFNL transcripts had markedly higher villus scores than E11⁺ cells lacking IFNL (**Figure 5D**). Together, these findings indicate that mature enterocytes are the predominant source of IFN-λ during E11 infection.

**Figure 5.**
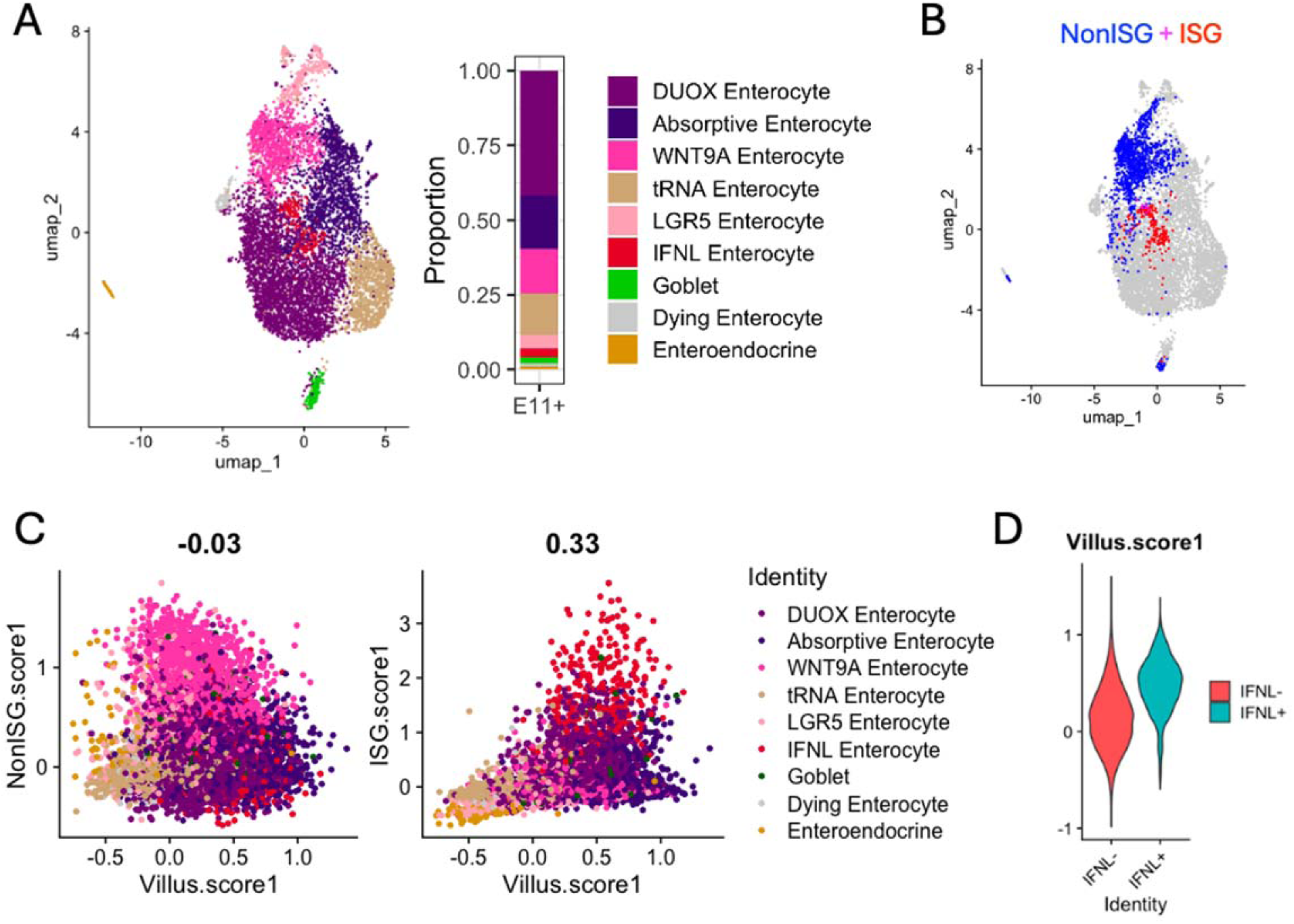
IFN-III signaling emanates from mature enterocytes. UMAP and bar plot showing the distribution and frequency of E11+ cells (A). UMAP with Non-ISG score (blue), ISG score (red), overlap (magenta), and neither (grey) shows the distribution of cells expressing the top 10 differentially expressed genes classified as ISGs or non-ISGs (B). Scatter plots show non-ISG score (left) and ISG score (right) with villus score for individual cells. Cells are colored by their associated cluster. The Pearson correlation coefficient is shown above the plot (C). Violin plot showing the distribution of villus scores for IFNL-(red) and IFNL+ (turquoise) cells (D).

### The IFN Response to Echovirus infection in Enterocytes Requires TLR3-TRIF Signaling

To determine which PRRs sense E11 in differentiated enteroids we used CRISPR-Cas9 to generate knockouts of the TLR3 adaptor protein TRIF, the RIG-I/MDA5 adaptor protein MAVS, and the transcription factor IRF3. Enteroids were dissociated into single-cell suspensions, transduced with lentiviruses carrying the respective gRNAs, and expanded as clonal populations. Clones were screened for successful gene knockout (KO) by detecting insertions/deletions (indels) at the CRISPR-Cas9 target site (**Figure S1-2**).

We next infected enteroid KO clones lacking TRIF, MAVS, or IRF3, as well as scrambled gRNA control clones, with E11 and harvested both RNA and supernatants 24 hours post-infection. Loss of TRIF or IRF3 completely abrogated the induction of IFN-λ1 and IFN-λ2, as well as the upregulation of the canonical ISG IFIT1 in response to infection (**Figure 6A–B**). In contrast, MAVS KO enteroids showed little to no impairment of IFN-λ or ISG induction, indicating that the RIG-I/MDA5–MAVS pathway is largely dispensable for E11 sensing in this system. MAVS immunoblots confirmed complete MAVS KO in our clones **(Figure S3**). Despite the differences in IFN responses, none of the knockouts including TRIF, IRF3, or MAVS altered the release of infectious E11 particles from infected enteroids (**Figure 6D**). This demonstrates that although dsRNA sensing through TLR3, TRIF, and IRF3 is essential for mounting an IFN-λ response to E11, the IFNs produced *de novo* were not sufficient to restrict viral replication or spread within enteroids.

**Figure 6.**
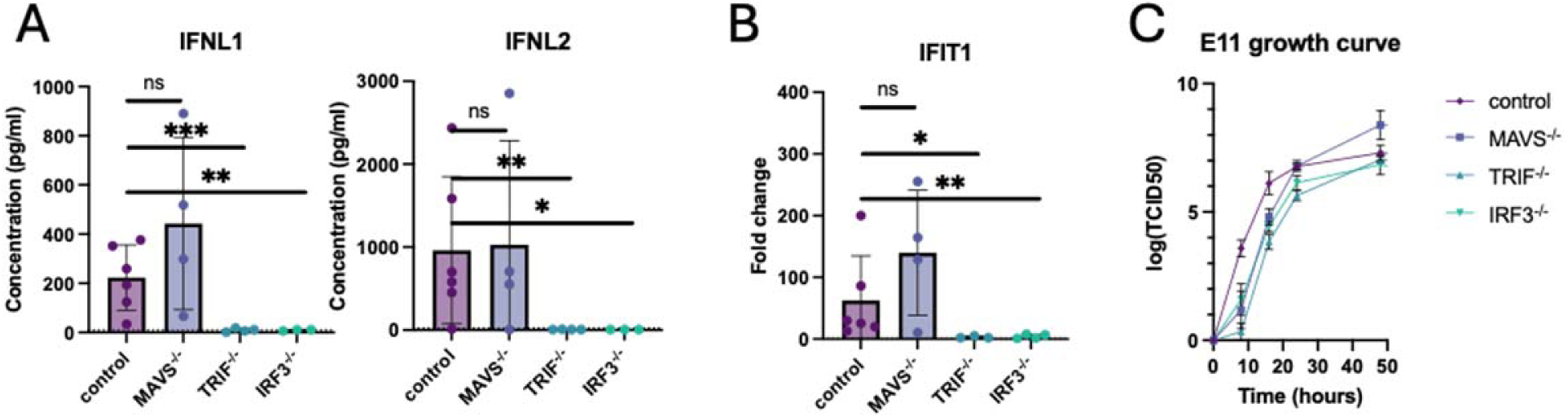
IFN-III induction by E11 requires TLR-TRIF signaling. Differentiated enteroid clones lacking MAVS, TRIF, or IRF3, or a scrambled gRNA control were infected with 1 pfu/cell E11. Supernatants and RNA were harvested 24 hours post-infection and used to measure IFNL1 and IFNL2 protein by Luminex (A) and IFIT1 transcript by qPCR (B). Differentiated enteroid clones were infected with 0.01 pfu/cell E11 and 100 µl of supernatant sampled 8, 16, 24, and 48 hours post-infection. Supernatants were used to measure infectious E11 by TCID50 assay (C). Mean, standard error, and individual replicates are shown for qPCR and Luminex experiments and the limit of detection (dotted line) is shown for Luminex experiments. Mean and standard error are shown for plaque assays. Mean values were log normalized and compared by one-way ANOVA with Dunnett’s multiple comparisons test. ns: not significant, *: p<0.05, **:p<0.005, ***:p<0.0005.

## Discussion

Our findings demonstrate that TLR3 is the primary driver of IFN-λ production in the intestinal epithelium during E11 infection, with mature enterocytes serving as the predominant source of this response. While both TLR3 and MDA5 have been implicated in anti-enteroviral defense at the organismal level, it has remained unclear which PRRs operate within the intestinal epithelium. By leveraging human-derived enteroids, our work identifies TLR3 as the key sensor of enteroviruses in this context and further reveals that enterocyte maturation enhances IFN-λ signaling capacity

Although *in vivo* studies demonstrate that IFN-III signaling is important for intestinal protection from enteroviruses, our data suggest that this response alone is insufficient to protect enteroids infected with E11 in vitro (25, 32). In our enteroid model, knockout of TRIF or IRF3 impaired IFN-λ production and ISG induction but had no detectable effect on E11 replication. This remained true even when we repeated infections with low multiplicities of E11, where no reproducible differences in viral genome levels or release of infectious virus were observed. Importantly, pretreatment of enteroids with exogenous IFN-λ restricted echovirus replication, consistent with the antiviral potential of this pathway. However, echoviruses are known to antagonize IFN-λ production and signaling as early as 2 hours post-infection, and complete their replication cycle within ∼6 hours (34, 43, 44). Thus, in the enteroid system, viral replication may simply outpace de novo IFN-λ responses, explaining why loss of TRIF or IRF3 does not alter viral yield despite their critical role in interferon induction.

Consistent with viral antagonism of IFN-λ, our scRNA-seq data revealed that only a small fraction of cells produced IFNL transcripts during E11 infection. At 16 hours post-infection, we detected E11 genomes in ∼40% of cells, yet only ∼4% upregulated IFNL, indicating that the vast majority of infected cells failed to mount an IFN response. This discrepancy underscores the ability of echoviruses to efficiently suppress or evade epithelial IFN-λ responses. The cellular basis for this heterogeneity remains unclear, but prior work in cell lines suggests that stochastic events upstream of interferon transcription, such as variability in PRR activation thresholds or chromatin accessibility, influence whether individual cells engage an antiviral program (45, 46). A likely explanation for the difference between our enteroid model and *in vivo* observations is the absence of immune and stromal cells *in vitro*. In the intact intestine, professional immune cells such as dendritic cells and macrophages, as well as stromal and endothelial populations, contribute additional layers of viral sensing, amplify IFN signaling, and provide paracrine cytokine support that may accelerate and broaden the epithelial antiviral response. Moreover, *in vivo* signaling networks, including cross-talk with type I and type II IFNs, can synergize with IFN-λ to enhance antiviral protection. In enteroids, by contrast, epithelial cells must generate the entire response autonomously, which may allow viral replication to outpace the limited and heterogeneous induction of IFN-λ.

Our transcriptional analyses of intestinal tissue and enteroids suggest that TLR3 and IFNLR1 expression increases as enterocytes mature, and that enteroids enriched for mature enterocytes mount stronger responses to both dsRNA and IFN-λ. Although prior studies have suggested that TLR3 and IFNLR1 are broadly expressed across epithelial cell types(12, 13, 47–49), there is evidence for heterogeneity as one report described higher TLR3 protein expression in differentiated enterocytes compared to crypt stem cells in the colon (50). Developmental regulation of these pathways may also be relevant. In mice, intestinal TLR3 expression increases during the first weeks of life, contributing to adult resistance to rotavirus infection (51), whereas neonatal protection relies more on constitutive ISG expression (52). These age-dependent changes in TLR3 and ISG expression coincide with the formation of intestinal villi and the emergence of mature enterocytes, providing a potential explanation for developmental differences in viral susceptibility (53).

It has been suggested that compartmentalization of IFN signaling is important to protect stem cells from the anti-proliferative effects of IFN signaling (54). While this concept has not been directly tested in ISCs, our findings are consistent with this model. By restricting IFN-λ production and responsiveness to mature villus enterocytes, the epithelium may establish a frontline antiviral barrier that shields ISCs in the crypts from both viral cytopathology and the growth-suppressive effects of interferon signaling.

In summary, our study defines TLR3–TRIF–IRF3 signaling in mature enterocytes as the primary pathway driving IFN-λ responses to E11 infection. While this response is robust at the transcriptional level, our enteroid model demonstrates that *de novo* IFN-λ production alone is insufficient to restrict echovirus replication *in vitro*, likely reflecting the combination of viral antagonism and the absence of immune and stromal amplifying circuits present *in vivo*. By linking IFN-λ induction to enterocyte maturation and villus development, our work provides a framework for understanding age-dependent susceptibility to enteroviruses and suggests that compartmentalization of IFN signaling may protect intestinal stem cells from its anti-proliferative effects. More broadly, these findings establish human enteroids as a powerful system to dissect epithelial-intrinsic antiviral defenses, while also highlighting the need for more complex models incorporating immune and stromal components to fully capture host-virus interactions at the intestinal barrier

## Materials & Methods

### Enteroid culture

Enteroids were derived from human tissue as previously described (34). Basal media contained 10mM HEPES (Gibco), 2 mM L-glutamine (Gibco), and 1x penicillin-streptomycin (Gibco) in Advanced DMEM/F12 (Gibco). Expansion, patterning, and differentiation media contained additional additives (Table S1). Enteroids were grown in expansion media renewed every 2-3 days. Enteroids were dissociated by resuspension in 500 µl of TrypLE (Gibco) for 5 min at 37 °C followed by addition of 500 µl of basal media. Enteroids were then mechanically disrupted, resuspensed in Matrigel, and plated as previously described (55). To pattern and differentiate enteroids, they were first mechanically disrupted in PBS and plated as previously described (6, 55). Patterning media was renewed every 1-3 days for 11 days followed by 3 days in either patterning or differentiation media.

### CRISPR-Cas9 KO

Lentivirus plasmids for packaging (pMD2.G), pseudotyping (psPAX2), and targeting (pLentiCRISPRv2) were ordered from VectorBuilder. Gene specific gRNAs were selected using VectorBuilder’s online tool. 5 µg of pMD2.G, 7.5 µg of psPAX2, and 10 µg of pLentiCRISPRv2 were transfected into a 175 cm^2^ flask of HEK 293T cells using Lipofectamine 3000 (Life Tech) according to the manufacturer’s instructions. Supernatants were harvested 48 and 72 hours later and centrifuged at 1,500 x g for 10 minutes to clear cell debris. Cleared supernatants were filtered through a 0.45 μm filter and then pelleted at 100,000 x g for 2 hours at 4 °C. The lentivirus pellet was resuspended in 1 ml of expansion medium containing 8 µg/ml polybrene and 10 µM Y-27632 (Roc inhibitor), aliquoted, and stored at −80 °C.

For transduction, enteroids were first dissociated to single cells by resuspension in 500 µl of TrypLE at 37°C for 15 min followed by mechanical disruption by forcefully pipetting up and down (55). Cells were resuspended in 300 µl of lentivirus, centrifuged at 600 x g for 1h, and incubated for 3 hours as previously described (56). Cells were then resuspended in 40 µl of Matrigel and plated as normal.

Enteroids were passaged once in 2 µg/ml puromycin to select for transduced cells, dissociated to single cells, and seeded at low density. After sufficient growth, domes were resuspended in Cell Recovery Solution (Corning) on ice for 30 minutes to remove Matrigel and single enteroids dissociated as above and passaged into 2 wells.

Enteroids from one well were cryopreserved and whole genomic DNA was purified from the second dome using a GenElute Mammalian gDNA Miniprep Kit (Sigma). Approximately 400 bp surrounding the gRNA target region was amplified by PCR using gene specific primers (Table S2), sequenced by sanger sequencing (Azenta), and ICE Analysis (Synthego) used to measure the proportion of insertion/deletions (indels). Protein extracts from MAVS KO clones were additionally probed by western blot to verify loss of protein expression (**Figure S1**).

### scRNA-seq

Enteroids were scraped into cold PBS, pelleted 5 min at 400 x g, resuspended in TrypLE, and incubated at 37°C for 15 min. Cells were then mechanically dissociated, diluted in basal media, pelleted 5 min at 400 xg, resuspended in DPBS with 1% BSA, and filtered through a 30 µm strainer to remove cell aggregates.

Libraries were prepared from ∼10,000 cells using 10x Genomics Single Cell 3’ reagent kits on a Chromium instrument (10x Genomics) by the Molecular Genomics Core at the Duke Molecular Physiology Institute. Sequencing was performed on a NovaSeq 6000 sequencer (Illumina) using a S2 flow cell which was predicted to yield around 62k reads/cell.

The CellRanger pipeline (10x Genomics) was used to filter bulk sequence data and map it to the human reference genome (GRCh38) or a custom genome with an added E11 genome sequence. Reads were then filtered to only include cells with 2-20% mitochondrial genes and 3000-10,000 genes. Datasets were then normalized and SCTransformed (57). Mitochondrial genes (percent.mt), number of reads (nCount_RNA), number of genes (nFeature_RNA), and the cell cycle scores (CC.difference) were regressed (58, 59). Biological replicates were then integrated with the IntegrateData function from Seurat v3 (60). Different treatment groups were then integrated using Harmony (61).

PCA, UMAP, and clustering were done with Seurat v3 (60). Adaptively-thresholded low rank approximation (ALRA) was used to impute values for low read transcripts and was used for subsequent analysis (62). Villus score was calculated using expression of 26 villus-associated genes (41). Differential expression analyses were carried out with DESeq2 (63).

### Immunofluorescence & Western blotting

For immunofluorescence, enteroids were stained in suspension in 1.5 ml tubes on a tube rotator as previously described (55). First, enteroids were resuspended in Cell Recovery Solution (Corning) on ice for 30 minutes to remove Matrigel and then resuspended in 4% PFA for 30 minutes at room temperature, washed in PBS, blocked/permeabilized in blocking buffer (2% goat serum in PBS) with 0.5% Tx-100 for 30 minutes, and washed in PBS. Enteroids were incubated with primary antibody diluted in blocking buffer overnight, washed in PBS, incubated with secondary antibody in blocking buffer for 1 hour at room temperature, washed with PBS, and mounted in VectaShield mounting media with DAPI (VectorLabs). 1:200 mouse anti-MUC2 clone 996/1 (Invitrogen), 1:200 rabbit anti-CHGA (Invitrogen), 1:200 mouse anti-Ki67 clone B56 (BD), 1:200 rabbit anti-Lysozyme C (DAKO), 1:400 phalloidin AF-633 (Invitrogen), 1:1000 anti-mouse AF-568 (Invitrogen), and 1:1000 anti-rabbit AF-488 (Invitrogen) were used for staining.

For western blots, 20 µg of protein lysates were separated on 4-20% polyacrylamide gradient gels (Biorad), transferred to nitrocellulose membranes, blocked with Intercept (PBS) blocking buffer (LICOR), stained overnight with 1:1000 polyclonal anti-MAVS (CST) in blocking buffer & 1 hour in 1:5000 anti-rabbit IRDye680 (LICOR) in blocking buffer, and scanned with an Azure 500 Imager (Azure Biosystems).

### E11 propagation, infection, & titration

E11 (Gregory) was obtained from the ATCC and propagated on HeLa cells (clone 7B) obtained from Jeff Bergelson, Children’s Hospital of Philadelphia, Philadelphia, PA. E11 was expanded on HeLa cells, concentrated by ultracentrifugation through a sucrose cushion, and titres determined by plaque assay on HeLa cells as described previously (64).

Enteroids were infected by resuspending domes in Cell Recovery Solution (Corning) on ice for 30 minutes to dissolve Matrigel. A fraction of enteroids were dissociated in TrypLE, as above, and cells counted on an automated cell counter (Biorad). Intact enteroids were pelleted at 200 x g for 5 minutes and then resuspended in 300 µl of basal media containing 0.01-1 pfu/cell E11 as indicated, added to a low-attachment 24-well plate, and rocked for 1 h at room temperature. Enteroids were washed once with basal media, pelleted as above, resuspended in 40 µl of Matrigel, and replated.

To measure E11 replication we used E11 containing Neutral Red and inactivated bound non-infectious virus by exposure to visible light as previously described (65, 66). After infection 100 µl of supernatant was sampled with replacement and infectious E11 measured using plaque assays or TCID50 as described previously (64, 67).

### qPCR and Luminex

RNA was harvested and reverse transcribed using an RNeasy mini prep kit (Qiagen) and an iScript cDNA synthesis kit (Bio-Rad) according to the manufacturers’ instructions. qPCR was carried out using an iScript One-Step RT-PCR kit with SYBR green (Bio-Rad) and gene-specific primer pairs (Table S2) according to the manufacturer’s instructions.

Cell culture supernatants were harvested and cytokines measured using IFNB, IL29, and IL28A singleplex beads from the Bio-Plex Pro Inflammation Panel 1 and a Bio-Plex Immunoassay System (Rio-Rad) according to the manufacturer’s instructions.

## Acknowledgments

We thank Joshua Hatterschide and Nora Guadalupe Ramirez (Duke University) for their advice while analyzing scRNA-seq data. We thank Liheng (Henry) Yang (Duke University) for his advice on intestinal organoid culture and differentiation. This project was supported by NIH R01 AI081759. The funder had no role in study design, data collection and analysis, decision to publish, or preparation of the manuscript. David Hare carried out investigation, formal analysis, and writing of the original draft. Carolyn Coyne provided resources, supervision, and editing of the original draft. Both were involved in conceptualization, methodology, software, and visualization.

---

**Figure S1.**
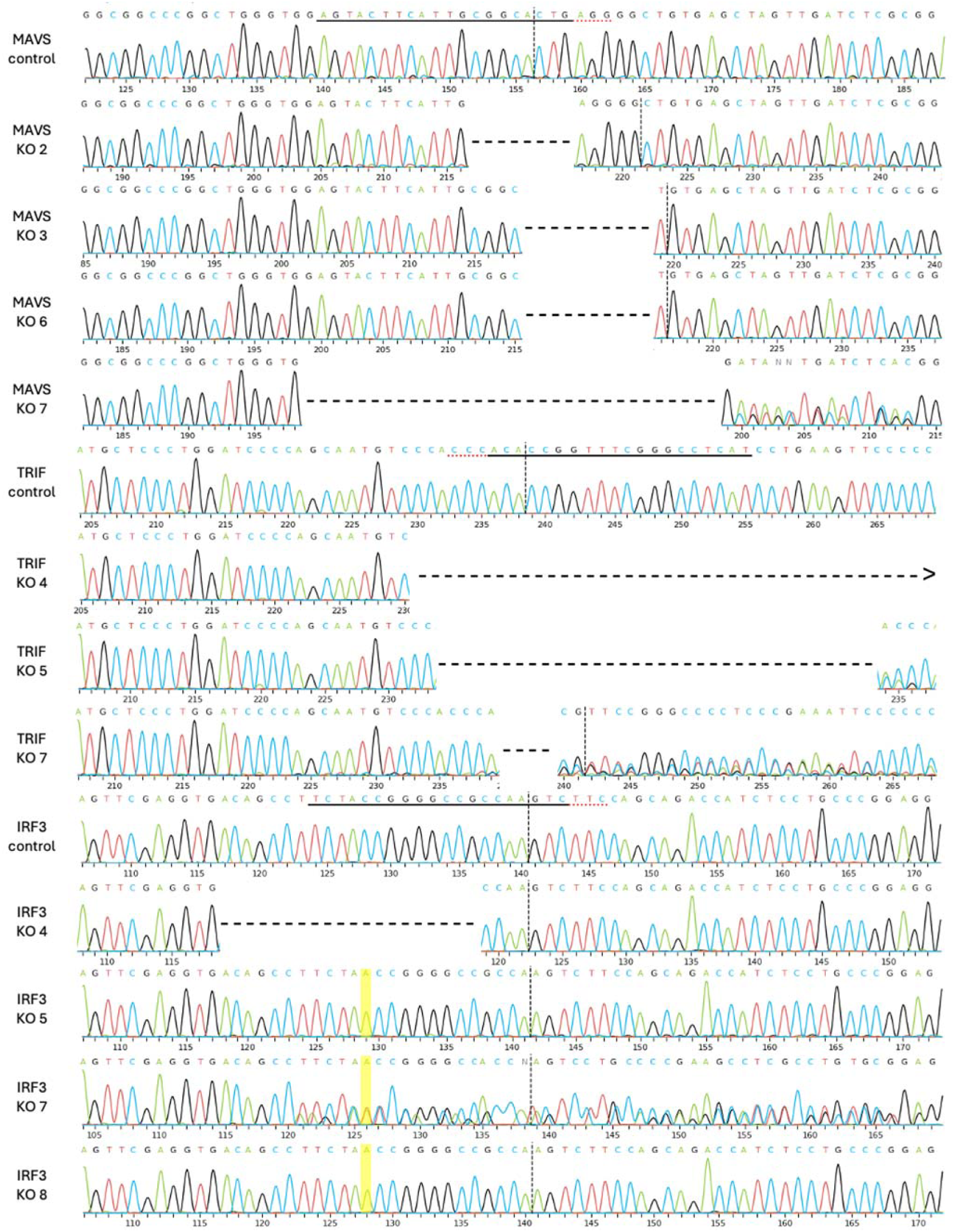
Sequence traces of KO clones. The sequencing trace is shown for control non-edited cells and each KO clone that was used. Deletions are shown by a dash for each deleted base while insertions are highlighted in yellow. The control sequence has the CRISPR target site underlined with a dotted line denoting the cut site. TRIF KO 4 had a deletion exceeding the sequence length.

**Figure S2.**
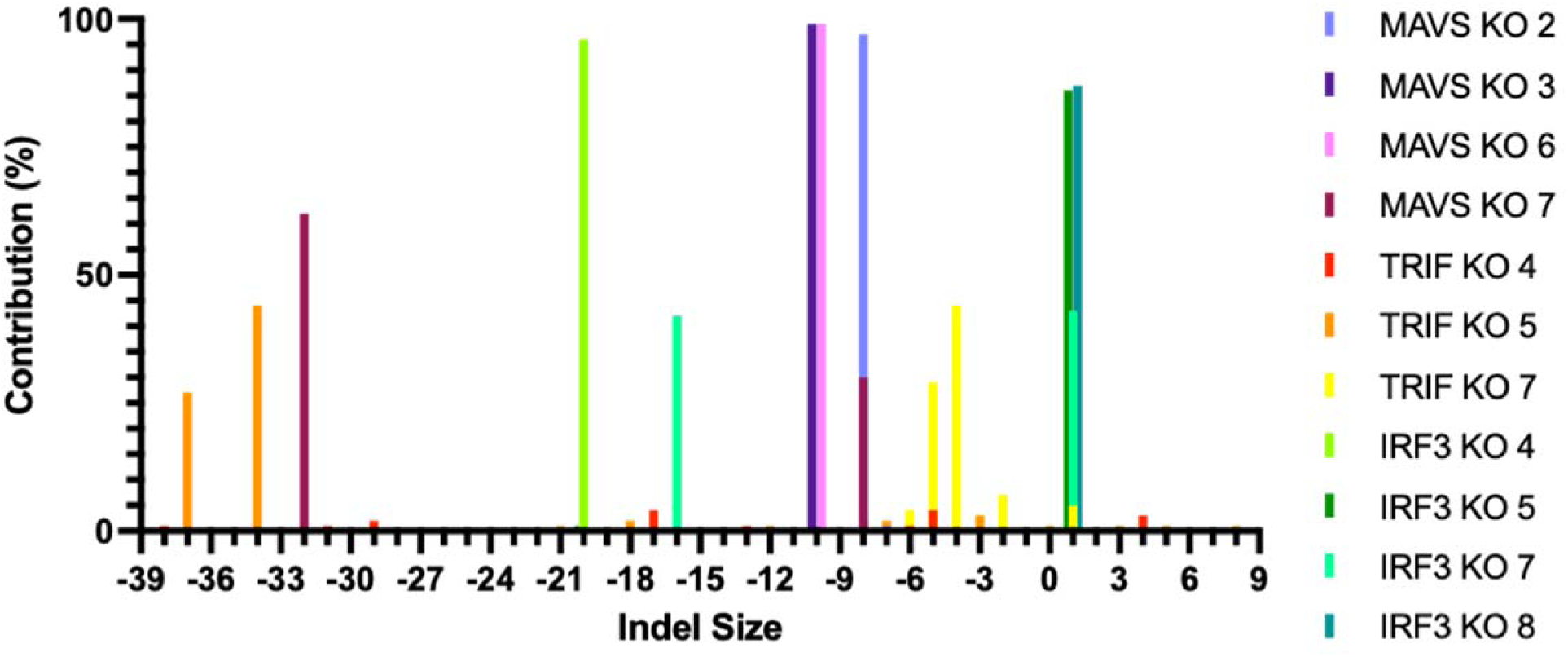
Indel contributions of KO clones. The size of indels and their relative contribution to the sequencing trace are plotted for each KO clone that was used. All indels with >1% contribution are shown. TRIF KO 4 had a large deletion confounding ICE analysis.

**Figure S3.**
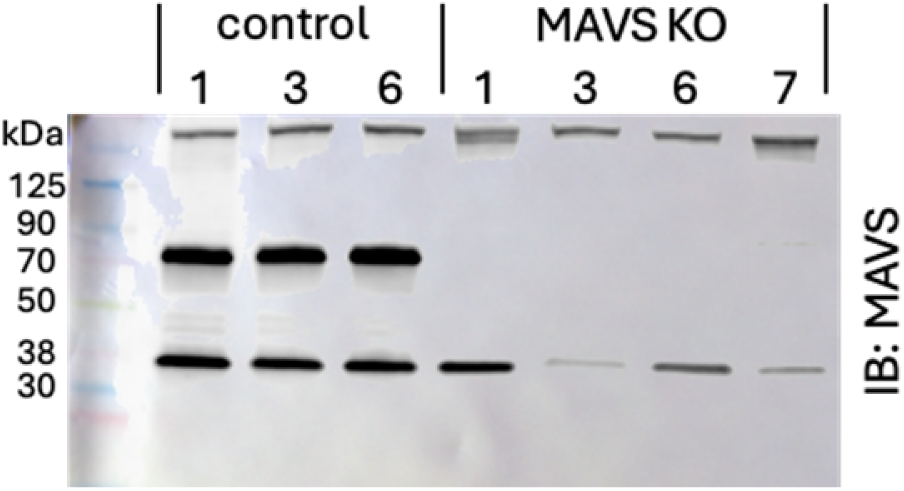
Western blot of MAVS KO clones. 20 µg of protein extracts from scrambled gRNA or MAVS KO clones were separated by SDS-PAGE and immunoblotted (IB) for the presence/absence of MAVS. MAVS (∼70 kDa) was detected in control clones above a non-specific band (∼35 kDa).

**Supplementary Table 1.**
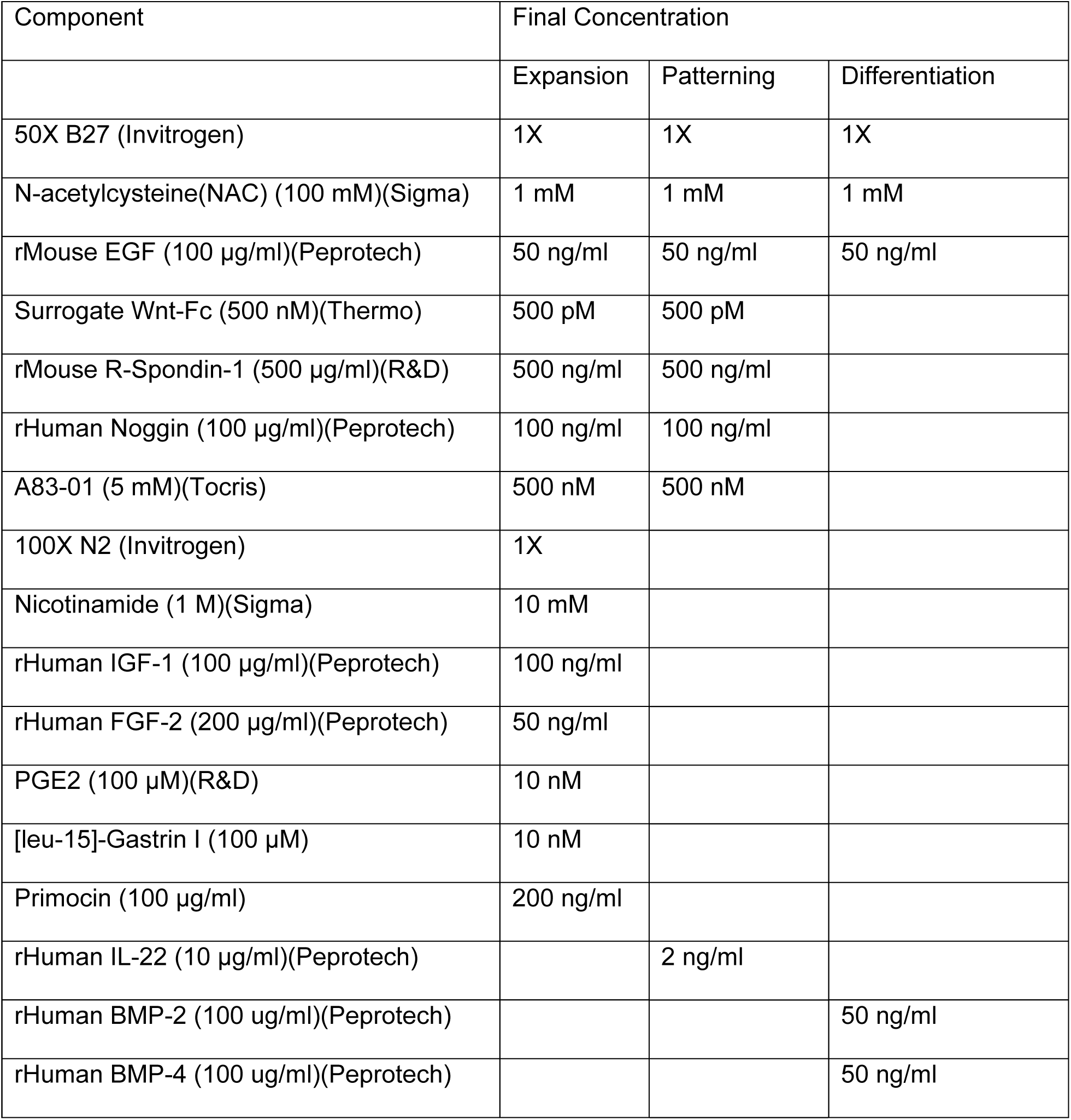
Enteroid media additives.

**Supplementary Table 2.**
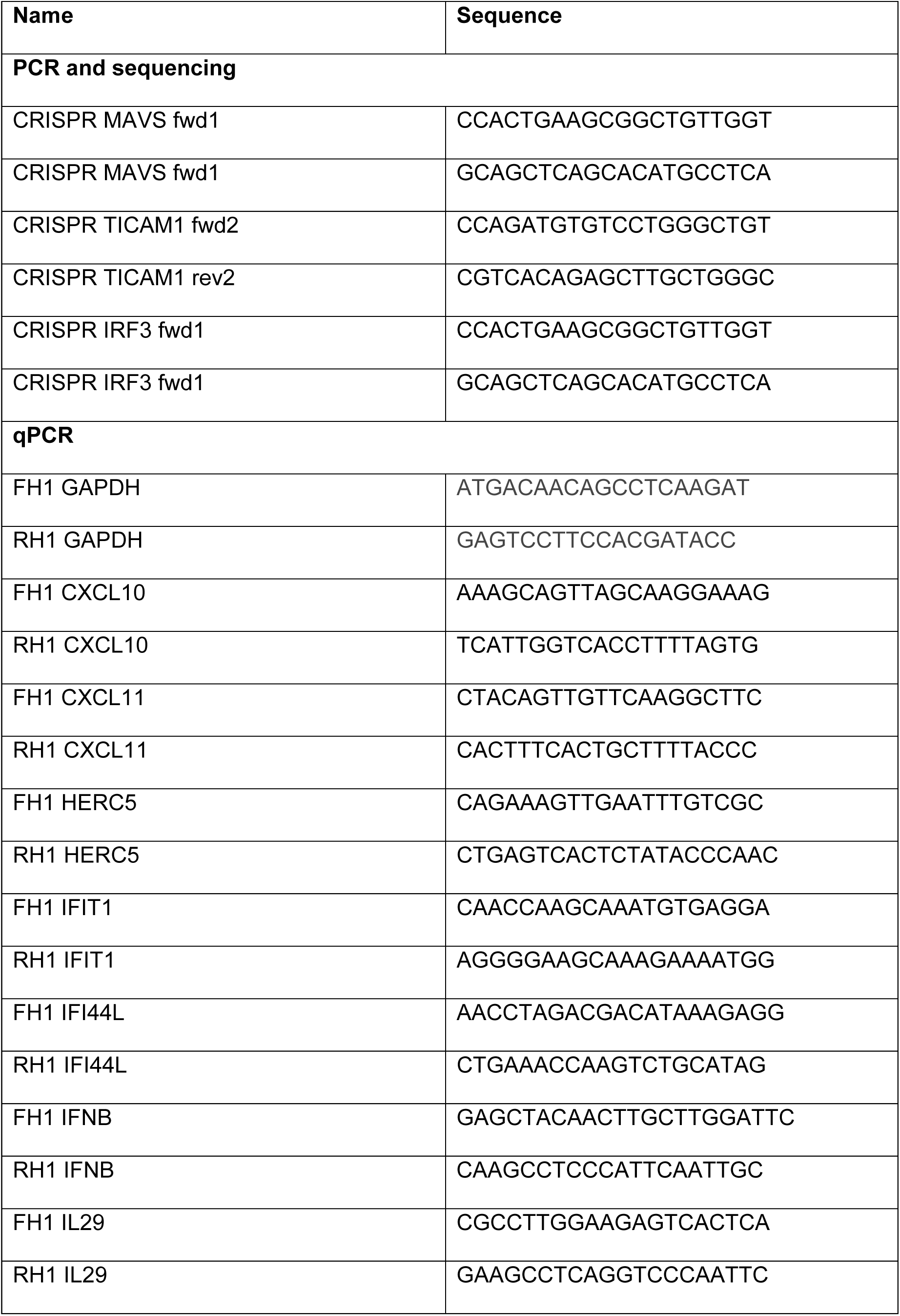
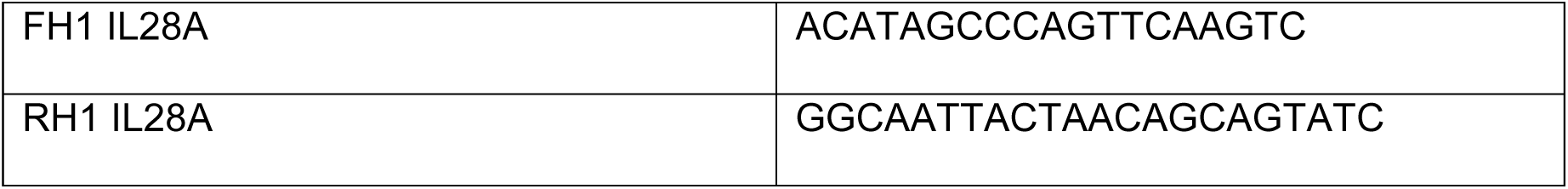
Primer sequences.

## References

1. Haston JC, Dixon TC. 2015. Nonpolio enterovirus infections in neonates. Pediatr Ann 44:e103–7.

2. Wells AI, Coyne CB. 2019. Enteroviruses: A Gut-Wrenching Game of Entry, Detection, and Evasion. Viruses 11.

3. van der Flier LG, Clevers H. 2009. Stem cells, self-renewal, and differentiation in the intestinal epithelium. Annu Rev Physiol 71:241–60.

4. Beumer J, Puschhof J, Yengej FY, Zhao L, Martinez-Silgado A, Blotenburg M, Begthel H, Boot C, van Oudenaarden A, Chen YG, Clevers H. 2022. BMP gradient along the intestinal villus axis controls zonated enterocyte and goblet cell states. Cell Rep 38:110438.

5. Sato T, Vries RG, Snippert HJ, van de Wetering M, Barker N, Stange DE, van Es JH, Abo A, Kujala P, Peters PJ, Clevers H. 2009. Single Lgr5 stem cells build crypt-villus structures in vitro without a mesenchymal niche. Nature 459:262–5.

6. He GW, Lin L, DeMartino J, Zheng X, Staliarova N, Dayton T, Begthel H, van de Wetering WJ, Bodewes E, van Zon J, Tans S, Lopez-Iglesias C, Peters PJ, Wu W, Kotlarz D, Klein C, Margaritis T, Holstege F, Clevers H. 2022. Optimized human intestinal organoid model reveals interleukin-22-dependency of paneth cell formation. Cell Stem Cell 29:1333–1345 e6.

7. Fujii M, Matano M, Toshimitsu K, Takano A, Mikami Y, Nishikori S, Sugimoto S, Sato T. 2018. Human Intestinal Organoids Maintain Self-Renewal Capacity and Cellular Diversity in Niche-Inspired Culture Condition. Cell Stem Cell 23:787–793 e6.

8. Lazear HM, Nice TJ, Diamond MS. 2015. Interferon-lambda: Immune Functions at Barrier Surfaces and Beyond. Immunity 43:15–28.

9. Kotenko SV, Gallagher G, Baurin VV, Lewis-Antes A, Shen M, Shah NK, Langer JA, Sheikh F, Dickensheets H, Donnelly RP. 2003. IFN-lambdas mediate antiviral protection through a distinct class II cytokine receptor complex. Nat Immunol 4:69–77.

10. Sheppard P, Kindsvogel W, Xu W, Henderson K, Schlutsmeyer S, Whitmore TE, Kuestner R, Garrigues U, Birks C, Roraback J, Ostrander C, Dong D, Shin J, Presnell S, Fox B, Haldeman B, Cooper E, Taft D, Gilbert T, Grant FJ, Tackett M, Krivan W, McKnight G, Clegg C, Foster D, Klucher KM. 2003. IL-28, IL-29 and their class II cytokine receptor IL-28R. Nat Immunol 4:63–8.

11. Novick D, Cohen B, Rubinstein M. 1994. The human interferon alpha/beta receptor: characterization and molecular cloning. Cell 77:391–400.

12. Zhou Z, Hamming OJ, Ank N, Paludan SR, Nielsen AL, Hartmann R. 2007. Type III interferon (IFN) induces a type I IFN-like response in a restricted subset of cells through signaling pathways involving both the Jak-STAT pathway and the mitogen-activated protein kinases. J Virol 81:7749–58.

13. Sommereyns C, Paul S, Staeheli P, Michiels T. 2008. IFN-lambda (IFN-lambda) is expressed in a tissue-dependent fashion and primarily acts on epithelial cells in vivo. PLoS Pathog 4:e1000017.

14. Doyle SE, Schreckhise H, Khuu-Duong K, Henderson K, Rosler R, Storey H, Yao L, Liu H, Barahmand-pour F, Sivakumar P, Chan C, Birks C, Foster D, Clegg CH, Wietzke-Braun P, Mihm S, Klucher KM. 2006. Interleukin-29 uses a type 1 interferon-like program to promote antiviral responses in human hepatocytes. Hepatology 44:896–906.

15. Bayer A, Lennemann NJ, Ouyang Y, Bramley JC, Morosky S, Marques ET, Jr., Cherry S, Sadovsky Y, Coyne CB. 2016. Type III Interferons Produced by Human Placental Trophoblasts Confer Protection against Zika Virus Infection. Cell Host Microbe 19:705–12.

16. Alexopoulou L, Holt AC, Medzhitov R, Flavell RA. 2001. Recognition of double-stranded RNA and activation of NF-kappaB by Toll-like receptor 3. Nature 413:732–8.

17. Feng Q, Hato SV, Langereis MA, Zoll J, Virgen-Slane R, Peisley A, Hur S, Semler BL, van Rij RP, van Kuppeveld FJ. 2012. MDA5 detects the double-stranded RNA replicative form in picornavirus-infected cells. Cell Rep 2:1187–96.

18. Dixit E, Boulant S, Zhang Y, Lee AS, Odendall C, Shum B, Hacohen N, Chen ZJ, Whelan SP, Fransen M, Nibert ML, Superti-Furga G, Kagan JC. 2010. Peroxisomes are signaling platforms for antiviral innate immunity. Cell 141:668–81.

19. Seth RB, Sun L, Ea CK, Chen ZJ. 2005. Identification and characterization of MAVS, a mitochondrial antiviral signaling protein that activates NF-kappaB and IRF 3. Cell 122:669–82.

20. Odendall C, Kagan JC. 2015. The unique regulation and functions of type III interferons in antiviral immunity. Curr Opin Virol 12:47–52.

21. Thomson SJ, Goh FG, Banks H, Krausgruber T, Kotenko SV, Foxwell BM, Udalova IA. 2009. The role of transposable elements in the regulation of IFN-lambda1 gene expression. Proc Natl Acad Sci U S A 106:11564–9.

22. Siegel R, Eskdale J, Gallagher G. 2011. Regulation of IFN-lambda1 promoter activity (IFN-lambda1/IL-29) in human airway epithelial cells. J Immunol 187:5636–44.

23. Tissari J, Siren J, Meri S, Julkunen I, Matikainen S. 2005. IFN-alpha enhances TLR3-mediated antiviral cytokine expression in human endothelial and epithelial cells by up-regulating TLR3 expression. J Immunol 174:4289–94.

24. Kang DC, Gopalkrishnan RV, Wu Q, Jankowsky E, Pyle AM, Fisher PB. 2002. mda-5: An interferon-inducible putative RNA helicase with double-stranded RNA-dependent ATPase activity and melanoma growth-suppressive properties. Proc Natl Acad Sci U S A 99:637–42.

25. Chen J, Jing H, Martin-Nalda A, Bastard P, Riviere JG, Liu Z, Colobran R, Lee D, Tung W, Manry J, Hasek M, Boucherit S, Lorenzo L, Rozenberg F, Aubart M, Abel L, Su HC, Soler Palacin P, Casanova JL, Zhang SY. 2021. Inborn errors of TLR3- or MDA5-dependent type I IFN immunity in children with enterovirus rhombencephalitis. J Exp Med 218.

26. Gorbea C, Makar KA, Pauschinger M, Pratt G, Bersola JL, Varela J, David RM, Banks L, Huang CH, Li H, Schultheiss HP, Towbin JA, Vallejo JG, Bowles NE. 2010. A role for Toll-like receptor 3 variants in host susceptibility to enteroviral myocarditis and dilated cardiomyopathy. J Biol Chem 285:23208–23.

27. Abe Y, Fujii K, Nagata N, Takeuchi O, Akira S, Oshiumi H, Matsumoto M, Seya T, Koike S. 2012. The toll-like receptor 3-mediated antiviral response is important for protection against poliovirus infection in poliovirus receptor transgenic mice. J Virol 86:185–94.

28. Yang J, Yang C, Guo N, Zhu K, Luo K, Zhang N, Zhao H, Cui Y, Chen L, Wang H, Gu J, Ge B, Qin CF, Leng Q. 2015. Type I Interferons Triggered through the Toll-Like Receptor 3-TRIF Pathway Control Coxsackievirus A16 Infection in Young Mice. J Virol 89:10860–7.

29. Negishi H, Osawa T, Ogami K, Ouyang X, Sakaguchi S, Koshiba R, Yanai H, Seko Y, Shitara H, Bishop K, Yonekawa H, Tamura T, Kaisho T, Taya C, Taniguchi T, Honda K. 2008. A critical link between Toll-like receptor 3 and type II interferon signaling pathways in antiviral innate immunity. Proc Natl Acad Sci U S A 105:20446–51.

30. Wang JP, Cerny A, Asher DR, Kurt-Jones EA, Bronson RT, Finberg RW. 2010. MDA5 and MAVS mediate type I interferon responses to coxsackie B virus. J Virol 84:254–60.

31. Richer MJ, Lavallee DJ, Shanina I, Horwitz MS. 2009. Toll-like receptor 3 signaling on macrophages is required for survival following coxsackievirus B4 infection. PLoS One 4:e4127.

32. Su R, Shereen MA, Zeng X, Liang Y, Li W, Ruan Z, Li Y, Liu W, Liu Y, Wu K, Luo Z, Wu J. 2020. The TLR3/IRF1/Type III IFN Axis Facilitates Antiviral Responses against Enterovirus Infections in the Intestine. mBio 11.

33. Drummond CG, Bolock AM, Ma C, Luke CJ, Good M, Coyne CB. 2017. Enteroviruses infect human enteroids and induce antiviral signaling in a cell lineage-specific manner. Proc Natl Acad Sci U S A 114:1672–1677.

34. Good C, Wells AI, Coyne CB. 2019. Type III interferon signaling restricts enterovirus 71 infection of goblet cells. Sci Adv 5:eaau4255.

35. Wells AI, Grimes KA, Coyne CB. 2022. Enterovirus Replication and Dissemination Are Differentially Controlled by Type I and III Interferons in the Gastrointestinal Tract. mBio 13:e0044322.

36. Burclaff J, Bliton RJ, Breau KA, Ok MT, Gomez-Martinez I, Ranek JS, Bhatt AP, Purvis JE, Woosley JT, Magness ST. 2022. A Proximal-to-Distal Survey of Healthy Adult Human Small Intestine and Colon Epithelium by Single-Cell Transcriptomics. Cell Mol Gastroenterol Hepatol 13:1554–1589.

37. Wang Y, Song W, Wang J, Wang T, Xiong X, Qi Z, Fu W, Yang X, Chen YG. 2020. Single-cell transcriptome analysis reveals differential nutrient absorption functions in human intestine. J Exp Med 217.

38. Triana S, Stanifer ML, Metz-Zumaran C, Shahraz M, Mukenhirn M, Kee C, Serger C, Koschny R, Ordonez-Rueda D, Paulsen M, Benes V, Boulant S, Alexandrov T. 2021. Single-cell transcriptomics reveals immune response of intestinal cell types to viral infection. Mol Syst Biol 17:e9833.

39. Moor AE, Harnik Y, Ben-Moshe S, Massasa EE, Rozenberg M, Eilam R, Bahar Halpern K, Itzkovitz S. 2018. Spatial Reconstruction of Single Enterocytes Uncovers Broad Zonation along the Intestinal Villus Axis. Cell 175:1156–1167 e15.

40. Beumer J, Artegiani B, Post Y, Reimann F, Gribble F, Nguyen TN, Zeng H, Van den Born M, Van Es JH, Clevers H. 2018. Enteroendocrine cells switch hormone expression along the crypt-to-villus BMP signalling gradient. Nat Cell Biol 20:909–916.

41. Elmentaite R, Ross ADB, Roberts K, James KR, Ortmann D, Gomes T, Nayak K, Tuck L, Pritchard S, Bayraktar OA, Heuschkel R, Vallier L, Teichmann SA, Zilbauer M. 2020. Single-Cell Sequencing of Developing Human Gut Reveals Transcriptional Links to Childhood Crohn’s Disease. Dev Cell 55:771–783 e5.

42. Rusinova I, Forster S, Yu S, Kannan A, Masse M, Cumming H, Chapman R, Hertzog PJ. 2013. Interferome v2.0: an updated database of annotated interferon-regulated genes. Nucleic Acids Res 41:D1040–6.

43. Mukherjee A, Morosky SA, Delorme-Axford E, Dybdahl-Sissoko N, Oberste MS, Wang T, Coyne CB. 2011. The coxsackievirus B 3C protease cleaves MAVS and TRIF to attenuate host type I interferon and apoptotic signaling. PLoS Pathog 7:e1001311.

44. Limpens RW, van der Schaar HM, Kumar D, Koster AJ, Snijder EJ, van Kuppeveld FJ, Barcena M. 2011. The transformation of enterovirus replication structures: a three-dimensional study of single- and double-membrane compartments. mBio 2.

45. Rand U, Rinas M, Schwerk J, Nohren G, Linnes M, Kroger A, Flossdorf M, Kaly-Kullai K, Hauser H, Hofer T, Koster M. 2012. Multi-layered stochasticity and paracrine signal propagation shape the type-I interferon response. Mol Syst Biol 8:584.

46. Zhao M, Zhang J, Phatnani H, Scheu S, Maniatis T. 2012. Stochastic expression of the interferon-beta gene. PLoS Biol 10:e1001249.

47. Baldridge MT, Lee S, Brown JJ, McAllister N, Urbanek K, Dermody TS, Nice TJ, Virgin HW. 2017. Expression of Ifnlr1 on Intestinal Epithelial Cells Is Critical to the Antiviral Effects of Interferon Lambda against Norovirus and Reovirus. J Virol 91.

48. Stanifer ML, Mukenhirn M, Muenchau S, Pervolaraki K, Kanaya T, Albrecht D, Odendall C, Hielscher T, Haucke V, Kagan JC, Bartfeld S, Ohno H, Boulant S. 2020. Asymmetric distribution of TLR3 leads to a polarized immune response in human intestinal epithelial cells. Nat Microbiol 5:181–191.

49. Price AE, Shamardani K, Lugo KA, Deguine J, Roberts AW, Lee BL, Barton GM. 2018. A Map of Toll-like Receptor Expression in the Intestinal Epithelium Reveals Distinct Spatial, Cell Type-Specific, and Temporal Patterns. Immunity 49:560–575 e6.

50. Furrie E, Macfarlane S, Thomson G, Macfarlane GT, Microbiology Gut Biology G, Tayside T, Tumour B. 2005. Toll-like receptors-2, -3 and -4 expression patterns on human colon and their regulation by mucosal-associated bacteria. Immunology 115:565–74.

51. Pott J, Stockinger S, Torow N, Smoczek A, Lindner C, McInerney G, Backhed F, Baumann U, Pabst O, Bleich A, Hornef MW. 2012. Age-dependent TLR3 expression of the intestinal epithelium contributes to rotavirus susceptibility. PLoS Pathog 8:e1002670.

52. Ramirez Reyes B, Madden S, Meyer KA, Bartsch B, Wright AP, Constant DA, Nice TJ. 2025. Homeostatic antiviral protection of the neonatal gut epithelium by interferon lambda. Cell Rep 44:115243.

53. Maier EA, Dusing MR, Wiginton DA. 2006. Temporal regulation of enhancer function in intestinal epithelium: a role for Onecut factors. J Biol Chem 281:32263–71.

54. Wu X, Dao Thi VL, Huang Y, Billerbeck E, Saha D, Hoffmann HH, Wang Y, Silva LAV, Sarbanes S, Sun T, Andrus L, Yu Y, Quirk C, Li M, MacDonald MR, Schneider WM, An X, Rosenberg BR, Rice CM. 2018. Intrinsic Immunity Shapes Viral Resistance of Stem Cells. Cell 172:423–438 e25.

55. Pleguezuelos-Manzano C, Puschhof J, van den Brink S, Geurts V, Beumer J, Clevers H. 2020. Establishment and Culture of Human Intestinal Organoids Derived from Adult Stem Cells. Curr Protoc Immunol 130:e106.

56. Van Lidth de Jeude JF, Vermeulen JL, Montenegro-Miranda PS, Van den Brink GR, Heijmans J. 2015. A protocol for lentiviral transduction and downstream analysis of intestinal organoids. J Vis Exp doi:10.3791/52531.

57. Hafemeister C, Satija R. 2019. Normalization and variance stabilization of single-cell RNA-seq data using regularized negative binomial regression. Genome Biol 20:296.

58. Hao Y, Hao S, Andersen-Nissen E, Mauck WM, 3rd, Zheng S, Butler A, Lee MJ, Wilk AJ, Darby C, Zager M, Hoffman P, Stoeckius M, Papalexi E, Mimitou EP, Jain J, Srivastava A, Stuart T, Fleming LM, Yeung B, Rogers AJ, McElrath JM, Blish CA, Gottardo R, Smibert P, Satija R. 2021. Integrated analysis of multimodal single-cell data. Cell 184:3573–3587 e29.

59. Satija R, Farrell JA, Gennert D, Schier AF, Regev A. 2015. Spatial reconstruction of single-cell gene expression data. Nat Biotechnol 33:495–502.

60. Stuart T, Butler A, Hoffman P, Hafemeister C, Papalexi E, Mauck WM, 3rd, Hao Y, Stoeckius M, Smibert P, Satija R. 2019. Comprehensive Integration of Single-Cell Data. Cell 177:1888–1902 e21.

61. Korsunsky I, Millard N, Fan J, Slowikowski K, Zhang F, Wei K, Baglaenko Y, Brenner M, Loh PR, Raychaudhuri S. 2019. Fast, sensitive and accurate integration of single-cell data with Harmony. Nat Methods 16:1289–1296.

62. Linderman GC, Zhao J, Roulis M, Bielecki P, Flavell RA, Nadler B, Kluger Y. 2022. Zero-preserving imputation of single-cell RNA-seq data. Nat Commun 13:192.

63. Love MI, Huber W, Anders S. 2014. Moderated estimation of fold change and dispersion for RNA-seq data with DESeq2. Genome Biol 15:550.

64. Morosky S, Lennemann NJ, Coyne CB. 2016. BPIFB6 Regulates Secretory Pathway Trafficking and Enterovirus Replication. J Virol 90:5098–107.

65. Harris KG, Morosky SA, Drummond CG, Patel M, Kim C, Stolz DB, Bergelson JM, Cherry S, Coyne CB. 2015. RIP3 Regulates Autophagy and Promotes Coxsackievirus B3 Infection of Intestinal Epithelial Cells. Cell Host Microbe 18:221–32.

66. Crowther D, Melnick JL. 1961. The incorporation of neutral red and acridine orange into developing poliovirus particles making them photosensitive. Virology 14:11–21.

67. Morosky S, Wells AI, Lemon K, Evans AS, Schamus S, Bakkenist CJ, Coyne CB. 2019. The neonatal Fc receptor is a pan-echovirus receptor. Proc Natl Acad Sci U S A 116:3758–3763.

